# First report of enhanced contents of nine macro- and micronutrients in gymnosperms via arbuscular mycorrhizal fungi

**DOI:** 10.1101/515031

**Authors:** Alicia Franco, Jesús Pérez-Moreno, Gabriela Sánchez, Carlos R. Cerdán, Juan J. Almaraz, Víctor M. Cetina, Alejandro Alarcón

## Abstract

Traditionally, it is thought that arbuscular mycorrhizae establish a mutualist symbiosis only with the roots of angiosperm plants. In this mutualism, fungi receive carbon from the plants, and angiosperms receive nutrients through the external mycelium of the arbuscular mycorrhizal fungi (AMF). However, the enhanced contents of macro- and micronutrients in gymnosperm plants, and therefore the mutualistic relationship, with AMF has not been reported so far. The present work evaluated whether arbuscular mycorrhizae were able to establish and enhance 9 nutrient contents in the neotropical Pinaceae species *Pinus greggii*. The tree seedlings were inoculated with three consortia of AMF isolated from an agricultural site, a forest of *Cupressus lusitanica* and a forest of *Pinus hartwegii*. The effect of AMF inoculation on plant growth and nutrient enhancement, in addition to colonization, was evaluated. There was evidence of enhancement of plant growth and 9 macro- and micronutrients in plants inoculated with the three evaluated consortia. After 7 months, the translocation was greater for Mg, Mn and Zn in plants inoculated with the consortium of AMF from pine forest. The presence of hyphae, vesicles and arbuscules was detected in the roots of the *Pinus greggii* plants inoculated with the AMF consortia. In addition to these positive effects, colonization of 10 to 15% and 20 to 38% was observed depending on the AMF consortia after 2 and 7 months, respectively. The presence of arbuscules which is the translocation structure among involved symbionts was also recorded; and photographed for the first time. In the present work, we report for the first time that arbuscular mycorrhiza affects the mobilization of N, P, K, Ca, Mg, Fe, Mn, Zn, Cu and B in gymnosperms, indicating that this mycorrhizal symbiosis is more complex than previously believed.

## Introduction

Under natural conditions, the majority of terrestrial plants form symbioses with mycorrhizae. Arbuscular mycorrhizal fungi (AMF) [1] are native to all terrestrial ecosystems and can be found in almost all soils [2]. These fungi are members of the phylum Glomeromycota [3,4] and are important components in the soil rhizosphere because they serve multiple functions in ecosystems, favour the growth of plants and facilitate the absorption of nutrients, including P, N and water [5,6]. AMF have been reported in most vascular plants, primarily in angiosperms. In contrast, it has been generally considered that gymnosperms are generally colonized by ectomycorrhizal fungi. However, there are some reports of colonization by AMF in Pinaceae [7, 8, 9]. For example, the presence of AMF vesicles in the roots of *Pseudotsuga menziesii* has been reported [15]. Additionally, several authors have reported AMF vesicles and hyphae in the roots of five other species of Pinaceae in the genera *Tsuga* [11, 12, 13], *Pinus* [14, 7, 15] and *Abies* [7]. Although the presence of AMF has been documented in six gymnosperm species, its functional importance in terms of nutrient enhancement has not been shown in this group of plants, which includes numerous species of importance to forests. In the present work, we studied the effect of the inoculation of three consortia of AMF on the growth and macronutrient (N, P, K, Ca and Mg) and micronutrient (Fe, Mn, Zn, Cu and B) content in the neotropical pine *Pinus greggii*. Mycorrhizal colonization was evaluated 2 and 7 months after inoculation.

## Materials and methods

### Biological materials and inoculum production

Rhizospheric soil was collected from three sites located in the community of San Pablo Ixayoc, Texcoco, state of Mexico over an altitude gradient of 2,650 m in the agricultural area, 2,700 m in the *Cupressus lusitanica* forest and 3,600 m in the *Pinus hartwegii* forest located on the western slope of the Tláloc Mountains, municipality of Texcoco. The rhizospheric soil of each site was used as an inoculum to propagate the AMF in each ecosystem. Pots with a 2 kg capacity were used, to which were added sterile river sand, 500 g of rhizospheric soil, and the seeds of corn and common grass (*Brachiaria decumbens*). These systems were used for the purpose of propagating the AMF present, with five pots per ecosystem over three months. Subsequently, the species present were identified and were termed the agricultural consortium (CSA), the cedar consortium (CSC) and the pine consortium (CSP).

### Inoculation of trees

We used *Pinus greggii* trees and obtained the seeds from a plantation in central Mexico in Toluca, state of Mexico. Prior to sowing, the seeds of *P. greggii* were soaked in distilled water for 24 hours to eliminate germination inhibiting compounds. The water was changed every seven hours to allow for the oxygenation of the embryos. The seeds were sterilized with 30% H_2_O_2_ for 20 minutes and rinsed with sterile distilled water under aseptic conditions. Once disinfected, the seeds were washed again for 15 minutes with sterile distilled water. Seeds were planted in a plastic container at a depth of 0.5 cm. Once germinated, the plants were transplanted into plastic tubes measuring 140 cm^3^ that contained the substrate, a mixture of river sand, crushed pine bark and forest soil at a 2:2:1 ratio. The substrate was sterilized with steam at 125 °C for 9 hours. Before transplanting, the tubes were filled at their base with a layer of sterilized tezontle to allow the flow of water during the experiment, and the rest was filled with sterilized substrate, including a layer of AMF of inoculum, according to the proposed treatments. A parallel set of plants was also set up without AMF inoculation according to the treatments. During the first 2 months after germination, a Captan solution was applied at a dose of 2 gL^-1^ of water every third day, for 20 days, followed by one application per week until the lignification of the stem occurred to avoid “damping off”, a disease commonly caused by *Phytophthora* sp., *Pythium* sp. and *Fusarium circinatum* [16]. The plants remained under greenhouse conditions for 210 days, at which time harvest was performed. The height, dry weight of the shoots and the roots, and mycorrhizal colonization were evaluated. A nutrient analysis was performed for N, P, K, Ca, Mg, Fe, Mn, Zn and B.

### Macro and micronutrients

Nutrient analyses were performed on the 10 plants used for the evaluation of dry weight. The N was determined by the semimicro-Kjeldahl method [17]. The total P was determined according to the method by Allen et al. [18]; K was extracted with ammonium acetate and measured by flame photometry. Ca, Mg, Fe, Cu, Mn, Zn and B were determined using atomic absorption spectrophotometry (Varian, Spectra-AA220).

### Mycorrhization

An adaptation of the clearing and staining method proposed by Phillips and Hayman was used [19]. The roots of *P. greggii* were placed in sterilisable plastic capsules in a beaker containing 10% KOH and incubated overnight. The following day, the samples were decanted and rinsed with running water. This process was repeated for five consecutive days. Next, H_2_O_2_ was applied for 1 hour, decanted and then rinsed with running water. Subsequently, 10% HCL was added for 1 hour and decanted, and then, 0.05% trypan blue dye was applied in lactoglycerol for 24 hours. The roots were cut into 1 cm long fragments that were then mounted on slides. Microscopic analysis was performed using light field optical microscopy to quantify the following AMF structures: hyphae, vesicles and arbuscules.

### Evaluation of variables

All *P. greggii* plants were harvested 7 months after sowing. At harvest, the height of the plants was evaluated, from the neck of the roots to the upper region of the apical bud. Each plant was extracted from the containers, and the root system was cut from the stem to the neck of the root. Subsequently, we performed a wash under running water to extract the largest amount of the root system. Sieves (0.180 and 0.085 mm) were used to reduce the loss of short roots. Next, to determine their dry weight, both the stems and the root system were dried at 80° C for 48 hours to a constant weight. This process was performed in 10 plants per treatment, as five plants were used to measure mycorrhizal colonization.

### Experimental design

The experimental design used four completely randomized treatments, including an uninoculated control and three treatments of plants inoculated with consortia of AMF isolated from agricultural soil, soil from a *Cupressus lusitanica* forest and soil from a *Pinus hartwegii* forest. These ecosystems were located in an altitudinal gradient ranging from 2,650 m in the agricultural area to 2,700 m in the *Cupressus* forest and 3,600 m in the P. *hartwegii* forest. Each of the four treatments, had 15 replicates; thus, the experiment consisted of a total of 60 experimental units, each consisting of a tree.

### Statistical analysis

For the variables of height, dry weight of the shoots and roots and nutritional content, an analysis of variance was performed, and a comparison of means was performed using Tukey’s test (P≤0.05) with the program Statistical Analysis System (SAS) [20]. The colonization data were transformed to their natural logarithms to meet the criteria of normality.

## Results

### Identification of AMF

The AMF morphotaxa of the three consortia studied were identified. The genera that predominated were *Glomus* and *Acaulospora*. In the agricultural area, *Cupressus lusitanica* forest and *Pinus hartwegii* forest, we found 16, 13 and 10 morphospecies of AMF, respectively (Fig 1). *Acaulospora scrobiculata* and *Archaeospora* sp. were found in all three sample areas. *Funneliformis mosseae* and *Scutellospora cerradensis* were found in both the agricultural area and in the *Cupressus lusitanica* forest.

**Figure 1.**
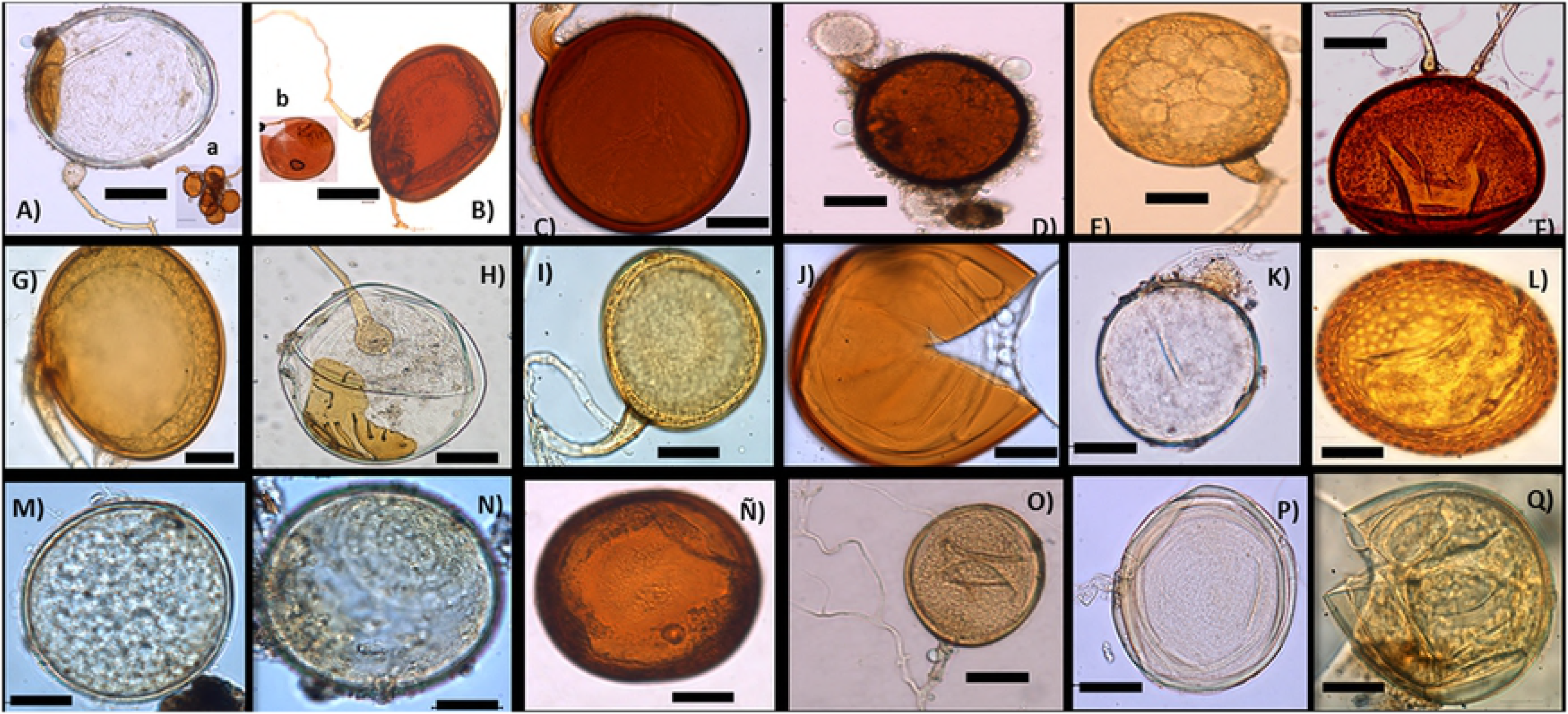
Species of arbuscular mycorrhizal fungi belonging to each of the three evaluated consortiums: fungi associated with the rhizosphere of the vegetation of the agricultural area (From A to F). A) *Scutellospora cerradensis* showing in (a) its helper cells; B) *Scutellospora pellucida* showing in (b) a broken spore; C) and D) *Septoglomus constrictum*; E) *Funneliformis mosseae*; F) *Gigaspora* sp.; Mycorrhizal fungi associated with the rhizosphere of the vegetation of *Cupressus lusitanica* (From G to L). G) *Claroideglomus etunicatum*; H) *Scutellospora cerradensis*; I) *Funneliformis mosseae*; J) *Acaulospora mellea*; K) *Archaeospora* sp.; L) *Acaulospora excavata.* Mycorrhizal fungi associated with the rhizosphere of *Pinus hartwegii* vegetation (From M to Q). M) and N) *Archaeospora* sp.; Ñ) *Acaulospora laevis*; O) *Glomus* sp.; P) *Paraglomus* sp.; Q) *Acaulospora* sp. Bar = 50 μm

### Plant growth

The *P. greggii* plants inoculated with the three AMF consortia showed increases in terms of growth and nutritional content. We observed an increase in height in mycorrhized plants compared to non-mycorrhized plants, which was independent of the evaluation time and the inoculated mycorrhizal consortium (Fig 2 a). Similar results were reported for *Abies lasiocarpa* when cultivated with AMF and a trap plant. We observed an increase in the dry weight of the shoots, especially 7 months after sowing, in inoculated plants compared to non-inoculated plants, regardless of the consortium (Fig 2 b). We observed an increase in the biomass of root dry weight 7 months after sowing in plants inoculated with AMF from the *Cupressus lusitanica* and *Pinus hartwegii* forests compared to those from the agricultural soil (Fig 2 c). In all three cases, the radical dry weight values were higher than those registered in non-inoculated plants (Fig 2 c). A trend similar to that recorded for the dry weight of the root was observed in the case of total dry weight (Fig 2 d).

**Figure 2.**
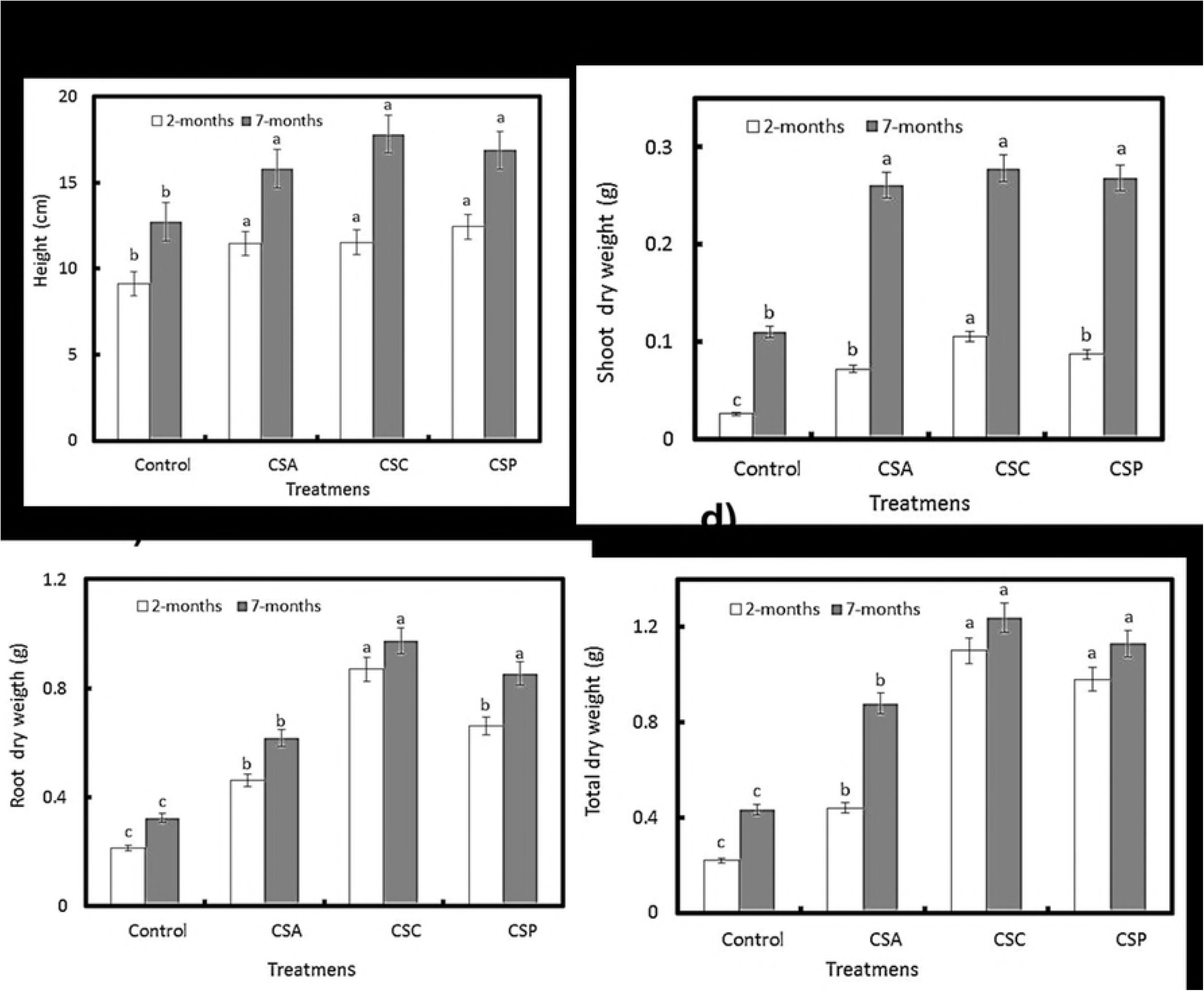
Growth of *Pinus greggii* a) Shoot height; b) Dry weight of the shoot; c) Dry weight of the root; d) Total dry weight. Inoculated with three consortia of arbuscular mycorrhizal fungi. CSA = consortium of agricultural land, CSC = Consortium of forest of *Cupressus lusitanica* and CSP = consortium of *Pinus hartwegii*. White bars = Plants 2 months old; Gray bars = 7-month-old plants. Equal letters above bars of the same color indicate equal values (Tukey, α = 0.05); n = 10 ± standard error of the mean.

### Mycorrhizal colonization

Differences were observed in terms of mycorrhizal colonization 2 and 7 months after sowing. Colonization values were 2.2 to 3.2 times higher 7 months after sowing, depending on the AMF consortium. The inoculated plants had mycorrhizal colonization values ranging from 20.5 to 35.5%, depending on the consortia (Fig 3). The presence of hyphae, vesicles and arbuscules was observed, as was the germination of AMF spores. In non-inoculated plants, no mycorrhizal colonization was observed (Fig 4).

**Figure 3.**
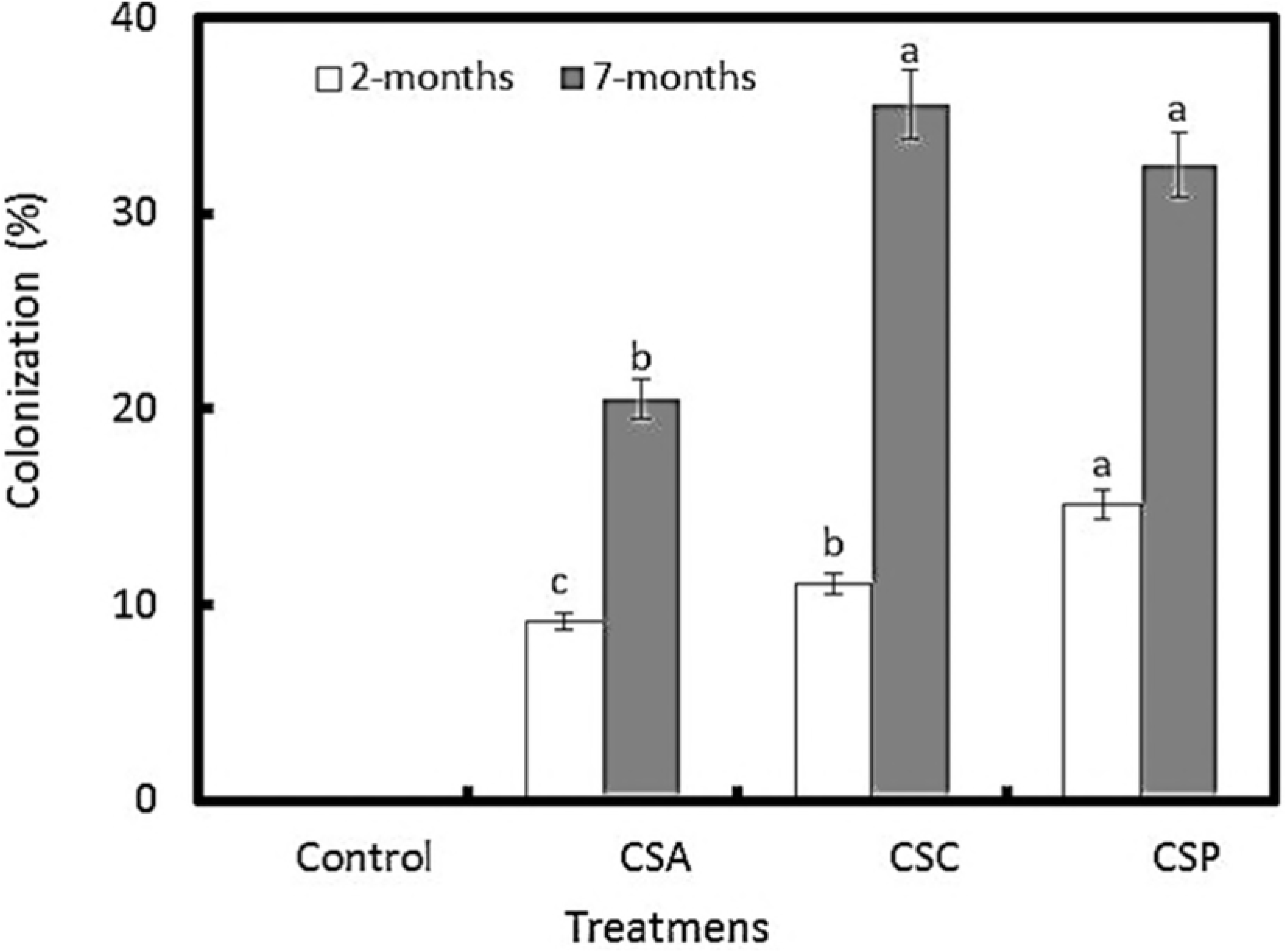
Colonization of *Pinus greggii* plants inoculated with three consortia of arbuscular mycorrhizal fungi. CSA = consortium of agricultural land, CSC = consortium of *Cupressus lusitanica* and CSP = consortium of *Pinus hartwegii*. White bars = Plants 2 months old; Gray bars = 7-month-old plants. Equal letters above bars of the same color are equal (Tukey, α = 0.05); n = 5 ± standard error of the mean.

**Figure 4.**
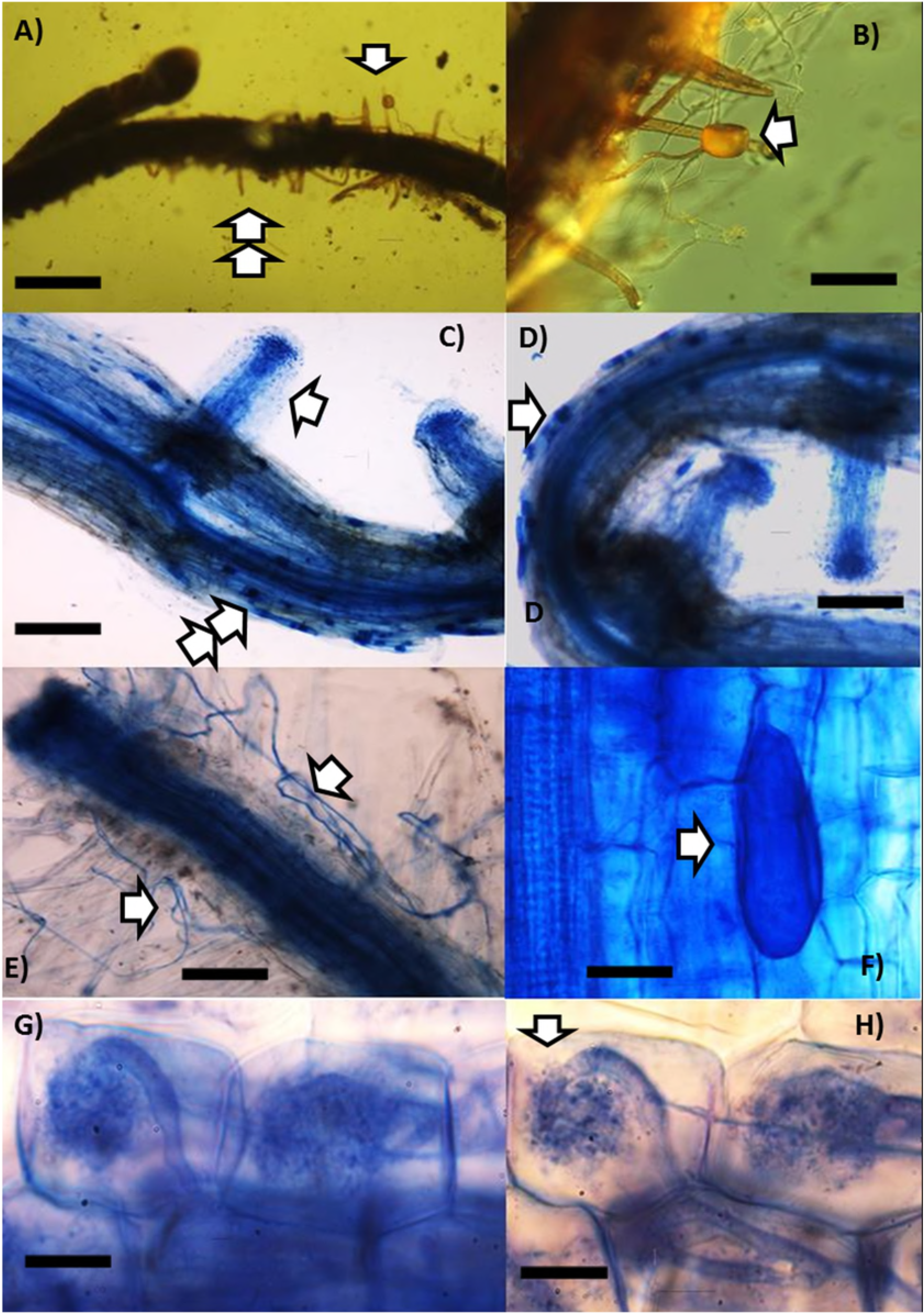
Colonization by arbuscular mycorrhizal fungi in *Pinus greggii* (Pg) seven months after inoculation, showing vesicles, external mycelium and arbuscules. A) Root of Pg showing radical hairs (double arrow) typical from pine roots; B) Close-up to a germinating *Glomus* sp. spore (arrow); C) Root of Pg showing abundant formation of vesicles (double arrow) and short feeding roots (arrow) typical from pine roots; D) Close-up to vesicles; E) Abundant extraradical hyphae of AMF; F) Close-up to a vesicle (arrow) within the cortical cells of Pg root from the *Cupresus* forest; G) and H) Arbuscules within the cortical cells of the Pg root of the AMF consortium of *Pinus hartwegii* forest. Bar = 30 μm.

### Nutritional content and transport

A higher content of the macronutrients N, P, K, Ca and Mg was observed in the shoots, in the roots and in total in inoculated plants compared to those not inoculated, regardless of the source of inoculum. A higher content was observed in plants inoculated with AMF from the *Cupressus lusitanica* forest (Table 1). A similar tendency was observed for the micronutrients Fe, Mn, Zn, Cu and B in the shoots, in the roots and in total in plants inoculated with AMF from the *Pinus hartwegii* forests (Table 2). The analysis of the nutrient content relationships in the shoots and roots of inoculated plants versus non-inoculated plants allowed us to determine the efficiency of nutrient transport resulting from the AMF. Based on these relationships, we observed the greatest transport for the macronutrients N, P, Mn and B in plants inoculated with AMF from *Pinus hartwegii* forests (Table 3). K, Ca and Mg in plants inoculated with agricultural soil; and Fe, Mn, Cu and B in plants inoculated with AMF from *Cupressus lusitanica* forests.

**Table 1.** Content of macronutrients (N, P, K, Ca and Mg) of the aerial part and root of *Pinus greggii* plants inoculated with three consortia of arbuscular mycorrhizal fungi (AMF), 210 days after sowing.

| Treatment ¶ | N | P | K | Ca | Mg |
| --- | --- | --- | --- | --- | --- |
| Aerial part |  |  |  |  |  |
| **Plants not inoculated** | 5.09 ± 0.23c | 0.19 ± 0.30b | 1.41 ± 0.04c | 1.22 ± 0.08c | **1.35 ± 0.05c** |
| Pi with CSA | 21.38 ± 0.65b | 0.79 ± 0.80a | 2.39 ± 0.21b | 5.98 ± 0.73b | **4.63 ± 0.08b** |
| Pi with CSP | 32.55 ± 0.36a | 0.89 ± 0.21a | 2.87 ± 0.28b | 6.10 ± 0.69a | **4.20 ± 0.07a** |
| Pi with CSC | 22.13 ± 0.71b | 0.99 ± 0.19a | 3.15 ± 0.36a | 5.75 ± 0.09b | **4.81 ± 0.06b** |
| Root |  |  |  |  |  |
| **Plants not inoculated** | 6.74 ± 0.21c | 0.23 ± 0.02c | 1.11 ± 0.03c | 1.80 ± 0.06b | **2.15 ± 0.04b** |
| Pi with CSA | 18.50 ± 0.58b | 0.67 ± 0.09b | 1.78 ± 0.08b | 5.24 ± 0.72a | **3.44 ± 0.05a** |
| Pi with CSP | 23.79 ± 0.61a | 0.64 ± 0.05b | 2.19 ± 0.19a | 5.64 ± 0.21a | **3.90 ± 0.03a** |
| Pi with CSC | 19.75 ± 0.51b | 0.76 ± 0.08a | 2.63 ± 0.24a | 5.32 ± 0.82a | **3.45 ± 0.02a** |
| Total |  |  |  |  |  |
| **Plants not inoculated** | 11.83±0.91c | 0.42±0.12b | 2.52±0.07c | 3.02±0.06b | **3.5±0.21b** |
| Pi with CSA | 39.88±0.99b | 1.46±0.06<sup>a</sup> | 4.17±0.12b | 11.22±0.82a | **8.07±0.63a** |
| Pi with CSP | 56.34±1.03<sup>a</sup> | 1.53±0.04a | 5.06±0.08a | 11.74±0.79a | **8.1±0.56a** |
| Pi with CSC | 41.88±0.97b | 1.75±0.21a | 5.78±0.09a | 11.07±0.81a | **8.26±0.81a** |

¶ CSA, consortium of agricultural land CSP; pine soil consortium; CSC; consortium of cedar floor. Pi = inoculated plants. Values are averages ± standard error of the mean, n = 10.

**Table 2.** Micronutrient content of the aerial part and root of *Pinus greggii* plants inoculated with three consortia of arbuscular mycorrhizal fungi (AMF), 210 days after sowing

| Treatment ¶ | Fe | Mn | Zn | Cu | B |
| --- | --- | --- | --- | --- | --- |
| Aerial part |  |  |  |  |  |
| **Plants not inoculated** | 0.12 ± 0.01d | 0.36 ± 0.21b | 0.28 ± 0.03c | 0.30 ± 0.02c | 0.42 ± 0.02b |
| Pi with CSA | 0.28 ± 0.02c | 0.71 ± 0.38a | 0.54 ± 0.02b | 0.69 ± 0.50b | 0.72 ± 0.51a |
| Pi with CSP | 0.49 ± 0.02a | 0.94 ± 0.45a | 0.85 ± 0.03a | 0.84 ± 0.71a | 0.90 ± 0.82a |
| Pi with CSC | 0.37 ± 0.02b | 0.87 ± 0.39a | 0.53 ± 0.03b | 0.79 ± 0.52b | 0.83 ± 0.52a |
| Root |  |  |  |  |  |
| **Plants not inoculated** | 0.18 ± 0.02c | 0.24 ± 0.02b | 0.36 ± 0.02c | 0.45 ± 0.03c | 0.55 ± 0.03b |
| Pi with CSA | 0.36 ± 0.03b | 0.55 ± 0.03a | 0.47 ± 0.02b | 0.64 ± 0.22b | 0.75 ± 0.22a |
| Pi with CSP | 0.69 ± 0.3a | 0.68 ± 0.023a | 0.73 ± 0.03a | 0.81 ± 0.71a | 0.84 ± 0.31a |
| Pi with CSC | 0.43 ± 0.02b | 0.55 ± 0.020a | 0.59 ± 0.05b | 0.73 ± 0.59b | 0.77 ± 0.26a |
| Total |  |  |  |  |  |
| **Plants not inoculated** | 0.30 ± 0.01c | 0.6 ± 0.02c | 0.64 ± 0.05c | 0.75 ± 0.03c | 0.97 ± 0.71b |
| Pi with CSA | 0.64 ± 0.3b | 1.26 ± 0.81b | 1.01 ± 0.32b | 1.33 ± 0.82b | 1.47 ± 0.82a |
| Pi with CSP | 1.18 ± 0.9<sup>a</sup> | 1.62 ± 0.92a | 1.58 ± 0.71a | 1.65 ± 0.91a | 1.74 ± 0.77a |
| Pi with CSC | 0.80 ± 0.05b | 1.42 ± 0.80a | 1.12 ± 0.36b | 1.52 ± 0.69a | 1.60 ± 0.58a |

¶ CSA, consortium of agricultural land CSP; pine soil consortium; CSC; consortium of cedar floor. Pi = inoculated plants. Values are averages ± standard error of the mean, n = 10.

**Table 3.** Aerial part relationships: macro root and micronutrients of *Pinus greggii* plants, 210 days after sowing inoculated with three consortia of mycorrhizal fungi.

| Treatment | N | P | K | Ca | Mg | Fe | Mn | Zn | Cu | B |
| --- | --- | --- | --- | --- | --- | --- | --- | --- | --- | --- |
| Pi with CSA | 1,15 | 1,18 | 1,34 | 1,14 | 1,34 | 0,77 | 1,29 | 1,14 | 1,07 | 1,04 |
| Pi with CSP | 1,36 | 1,39 | 1,31 | 1,08 | 1,07 | 0,71 | 1,38 | 1,16 | 1,03 | 1,07 |
| Pi with CSC | 1,12 | 1,30 | 1,19 | 1,08 | 1,21 | 0,86 | 1,58 | 1,11 | 1,08 | 1,07 |
| Psi | 0,75 | 0,82 | 0,82 | 0,67 | 0,62 | 0,66 | 0,15 | 0,77 | 0,66 | 0,76 |
| Pi with CSA:<br>Psi | 1,53 | 1,43 | **1,63** | **1,70** | **2,16** | 1,16 | 8,6 | 1,48 | 1,62 | 1,36 |
| Pi with CSP:<br>Psi | **1,81** | **1,69** | 1,59 | 1,61 | 1,72 | 1,07 | **9,2** | 1,50 | 1,56 | **1,40** |
| Pi with CSC:<br>Psi | 1,49 | 1,58 | 1,45 | 1,61 | 1,95 | **1,30** | **10,5** | 1,44 | **1,63** | **1,40** |

¶ CSA, consortium of agricultural land CSP; pine soil consortium; CSC; consortium of cedar floor. Pi = inoculated plants. Values are averages ± standard error of the mean, n = 10.

## Discussion

### Plant growth

Mycorrhizal fungi contribute to the growth and development of vascular plants. Multiple studies have shown that mycorrhizae also protect plants against soil pathogens. The increase in plant growth in terms of height and biomass after inoculation with AMF has been widely documented in angiosperms, but not in gymnosperms [1]. In the present work, *Pinus greggii* plants inoculated with AMF had increased height and greater shoot and root biomass compared to the non-inoculated plants.

### Mycorrhizal colonization

AMF are characterized by intra- and intercellular growth in the root cortex and by the formation of hyphae and external hyphae. In the present work, colonization of 20 to 40% was observed in the *Pinus greggii* plants with the AMF consortia in addition to the presence of hyphae, vesicles and arbuscules. There are reports of AMF colonization in gymnosperms. Several authors [7, 11–15] have reported the presence of AMF hyphae and vesicles in the roots of *Pseudotsuga menziesii, Tsuga, Abies* and *P. muricata*. These reports indicate that although ectomycorrhizal symbioses are predominant in the structure and function of gymnosperm roots, AMF are present in the roots of these trees as well.

### Nutrient mobilization

In the present work, the enhancement of macro- and micronutrients contents via AMF in gymnosperms is reported for the first time. Inoculation with all three AMF consortia in the neotropical pine *P. greggii* showed a beneficial effect in terms of growth, colonization and content of macro- and micronutrients in comparison with non-inoculated plants. Although nutrient mobilization by AMF has been extensively demonstrated in angiosperms, there are no reports of conspicuous nutrient contents enhancement in gymnosperms. This work reports the mobilization of N, K, Ca, Mg, Fe, Mn, Zn, Cu and B within the shoots and roots of gymnosperms. Previously, only two studies have reported a higher P content in the gymnosperm *Pseudotsuga menziesii* [11, 12].

In the present work, Mg, Mn and Zn were mobilized in the shoots of plants inoculated with AMF. Mn contributes to the functioning of various biological processes, including photosynthesis, via the synthesis of chlorophyll, respiration and the assimilation of nitrogen. Mn also participates in the formation of chloroplasts, activates the growth of plants, promotes cellular lengthening in the roots and confers resistance to pathogens. The transfer of Mn in angiosperms inoculated with mycorrhizal fungi has been reported. For example, Bethlenfalvay and Franson [21] recorded high concentrations of Mn in the shoots of barley plants (*Hordeum vulgare*). The same authors reported increases in Mn and increased growth in wheat plants (*Triticum durum*) inoculated with the AMF *Glomus monosporum*. On the other hand Arines et al. [22] found that in red clover (*Trifolium pratense*), the total Mn transfer increased in plants inoculated with the mycorrhizal fungi *Gigaspora aurigloba* and *Glomus tenue* compared with non-inoculated plants. In the present work, the difference in the Mn content between mycorrhized and non-mycorrhized trees was greater in the roots than in the shoots, possibly because the mycorrhizae altered the spatial distribution of this nutrient. The lower absorption of Mn by mycorrhized plants can be explained by the existence of a fungal mechanism that controls the absorption of Mn or by the effect of the fungi on the rhizosphere and surrounding soil. Previously, a decrease in Mn toxicity in the presence of AMF in soybeans has been documented.

Angiosperms colonized by AMF are often more resistant to excess Mn than plants not colonized by this fungus. Mg is an essential nutrient for plants and is critical for a wide range of functions. Mg is involved in photosynthesis and is a basic component of chlorophyll. Xiao et al. [23] observed greater biomass in the roots and shoots in orange plants (*Poncirus trifoliata*) and concluded that inoculation with the mycorrhizal fungus *Funneliformis mosseae* had positive effects on the growth and physiology under Mg-deficient conditions. Hassan Zare-Maivan et al. [24] observed that plants inoculated with mycorrhizae had significantly higher content of dry and fresh root weight and chlorophyll content than plants without mycorrhizae. Mycorrhizal colonization increased Mg uptake but decreased K uptake. Xiao et al. [23] suggested that the mycorrhizal fungus *Glomus versiforme* can improve the growth and distribution of Mg in orange seedlings grown in soil low in Mg. These authors reported that the concentrations of Mg in the shoots and roots, biomass yield and chlorophyll content increased with the inoculation of three species of mycorrhizal fungi, especially *Glomus versiforme*.

In the present work, significant nutritional transport of 10 macro- and micronutrients was observed, primarily Mg and Mn in the shoots of plants inoculated with the AMF consortium from the *Pinus hartwegii* forest.

## Conclusions

This work demonstrates for the first time the functional importance of AMF in terms of growth and nutrient enhancement contents in gymnosperms. AMF allowed for the mobilization of nine nutrients, primarily Mg, Mn and Zn, to the roots and shoots of the gymnosperm *Pinus greggii*. Plants of *P. greggii* inoculated with AMF produced more biomass than non-inoculated plants. The total colonization of *P. greggii* varied depending on the source of inoculum 7 months after inoculation. Greater colonization was observed in *Pinus* plants inoculated with the mycorrhizal consortia from *Cupressus* forests. Additionally, this is the first study that illustrates the formation of arbuscules by arbuscular mycorrhizal fungi in gymnosperms. The presence of arbuscules, which we documented photographically for the first time in gymnosperm plants, shows that P. *greggii* establishes a functional mutualist symbiosis with the AMF, as the exchange of nutrients occurs in this structure. These results indicate that *Pinus greggii* improves its nutritional status in the early stages of its development by associating with AMF; thus, inoculation with these fungi should be considered if reforestation activities of pine forests are desired.

## Aknowledgements

The first author thanks a PhD scholarship from CONACyT. The author of correspondance acknowledge CONACyT 2018-000007-01EXTV and COMECyT for the financial support to carry out an international sabbatical stay in the Kunming Institute of Botany, Chinese Academy of Sciences in Kunming, China.

